# Exploring the potential value of aloe active components in inhibiting non-small cell lung cancer through network pharmacology

**DOI:** 10.64898/2025.12.02.691806

**Authors:** Chen Shen, Mengxi Zhu, Jia Li, Yi Wang, Shu-Feng Zhou

## Abstract

Non-small cell lung cancer (NSCLC) remains the leading cause of cancer-related mortality worldwide, and its management is challenged by profound molecular heterogeneity and inevitable resistance to targeted therapies. Natural products offer a valuable reservoir for anticancer drug discovery, yet the therapeutic potential of Aloe vera—a medicinal plant rich in structurally diverse bioactive compounds—remains insufficiently characterised in the context of NSCLC. Here, we employed an integrated strategy combining network pharmacology, protein–protein interaction analysis, Gene Ontology and KEGG enrichment, molecular docking, and 100-ns molecular dynamics simulations to comprehensively elucidate the multi-target and multi-pathway regulatory mechanisms of Aloe vera active constituents against NSCLC. We identified 369 aloe-derived targets and 5519 NSCLC-related genes, with 268 overlapping targets enriched in pathways involving EGFR signalling, PI3K/AKT/mTOR activation, p53 regulation, inflammation, apoptosis, and drug-resistance networks. Core hub genes included TP53, EGFR, AKT1, and STAT3. Molecular docking demonstrated strong spontaneous binding between key aloe compounds (such as aloe-emodin, aloeresin C, quercetin and sitosterols) and wild-type EGFR, mutant EGFR T790M/C797S, AKT1, and p53. Molecular dynamics simulations confirmed stable ligand–protein interactions, favourable hydrogen-bonding networks, and energetically favourable binding free energies. Collectively, our findings reveal a systematic, multi-level regulatory landscape through which Aloe vera may exert anti-NSCLC effects and highlight several candidate molecules with potential to enhance therapeutic efficacy or overcome EGFR-TKI resistance. This study provides a solid theoretical framework for subsequent experimental validation and the development of novel aloe-derived anticancer agents with low toxicity.

## 1. Introduction

Lung cancer remains one of the most formidable global health challenges, displaying persistently high incidence and mortality across both developed and developing countries [1]. According to the most recent GLOBOCAN 2022 estimates, lung cancer accounted for approximately 2.48 million new diagnoses and 1.82 million deaths worldwide, representing the leading cause of cancer-related mortality [2]. Although clinical management of lung cancer has undergone profound transformation over the past two decades—particularly with the advent of molecular-targeted therapies and immune checkpoint inhibitors—survival benefits remain limited for a substantial proportion of patients [3]. Among the subtypes of lung cancer, non-small cell lung cancer (NSCLC) accounts for 80–85% of cases and is characterized by complex genomic alterations, heterogeneous tumor evolution, and high drug resistance, all of which contribute to poor prognosis and limited long-term survival [4].

NSCLC originates predominantly from bronchial or alveolar epithelial cells and is associated with diverse risk factors. Unlike small cell lung cancer (SCLC), which is strongly linked to long-term smoking and neuroendocrine transformation [5, 6], NSCLC may also arise in non-smokers. Contributing risk factors include germline susceptibility, chronic lung diseases, environmental pollutants, and occupational exposures [7, 8]. Recent multi-omics profiling has further revealed that NSCLC is not a singular disease entity; instead, it comprises multiple molecular subtypes defined by mutations in EGFR, KRAS, ALK, TP53, BRAF, MET, and others [9]. These molecular alterations influence tumor progression, immune evasion, metastatic potential, and treatment responsiveness.

Despite major progress in targeted therapy, resistance remains inevitable. First- and second-generation EGFR tyrosine kinase inhibitors (EGFR-TKIs) such as gefitinib, erlotinib, afatinib, and dacomitinib provide clinical benefits but commonly induce acquired resistance through EGFR T790M mutation, bypass pathway activation, histological transformation, or downstream signaling rewiring. Even third-generation TKIs such as osimertinib, although effective against T790M, ultimately fail due to the emergence of C797S mutation, MET amplification, epithelial–mesenchymal transition (EMT), or activation of the PI3K/AKT and RAS/MAPK pathways [10]. In addition, approximately 5% of EGFR-mutant NSCLC patients develop SCLC transformation, which leads to high proliferative capacity and profound drug resistance [11, 12]. Therefore, identifying new low-toxicity compounds with multi-target activity that can overcome or delay TKI resistance represents an urgent unmet clinical need.

Natural products derived from medicinal plants have long served as foundational sources for modern drug discovery. More than 60% of clinical anticancer drugs are directly or indirectly derived from natural compounds [13]. Traditional Chinese medicine (TCM) in particular contains a rich diversity of biologically active small molecules with multi-target, multi-pathway therapeutic potential. However, their application remains limited by unclear molecular mechanisms, complex phytochemical composition, and lack of systematic target verification [14]. Aloe vera (*Aloe vera (L.) Burm. f.*) is a perennial medicinal plant widely used for its anti-inflammatory, antibacterial, antioxidant, wound-healing, immunomodulatory, and metabolic regulatory properties [15, 16]. Its leaves contain more than 200 structurally diverse compounds, including polysaccharides (acemannan), anthraquinones (aloe-emodin), chromones (aloesin, aloeresin C), phytosterols (β-sitosterol), flavonoids (quercetin), vitamins, terpenoids, amino acids, lignin, and saponins. These molecules exert extensive biological effects, many of which have demonstrated anti-tumor activity through the induction of apoptosis, cell cycle arrest, anti-angiogenesis, inhibition of metastasis, and modulation of tumor immunity [17, 18].

Increasing evidence suggests that aloe-derived compounds possess substantial anticancer potential in colorectal, breast, liver, and skin cancers [19, 20]. For example, aloe-emodin induces mitochondrial-dependent apoptosis and disrupts redox homeostasis; aloeresin C exhibits potent protein kinase inhibition; and acemannan activates innate and adaptive immune responses. However, studies focusing on the anti-NSCLC potential of Aloe vera remain extremely limited. Preliminary research reported that aloe gel extract shows cytotoxic effects in lung cancer cell lines [21], yet the underlying molecular basis remains poorly understood.

Network pharmacology has recently emerged as a powerful, system-level methodology integrating bioinformatics, cheminformatics, multi-omics data, and computational modeling to elucidate the complex interactions between natural products and disease networks [22]. Unlike conventional “one drug–one target” paradigms, network pharmacology reveals multi-target synergy and topological regulation within disease pathways. When combined with molecular docking and molecular dynamics simulation (MDS), this integrative framework enables precise prediction of ligand–protein binding modes, complex stability, and thermodynamic behavior at atomistic resolution.

In this study, we conducted a comprehensive network pharmacology analysis to elucidate the multi-target mechanisms of Aloe vera active components against NSCLC. We systematically screened aloe-derived functional compounds, predicted and validated potential molecular targets, and constructed interaction networks that identify key regulatory hubs. We performed GO and KEGG enrichment analyses to map aloe–NSCLC-associated biological processes and signaling pathways. In addition, using molecular docking and 100-ns molecular dynamics simulations, we evaluated the binding affinities and dynamic stability between major aloe compounds and critical NSCLC targets including p53, wild-type EGFR, mutant EGFR T790M/C797S, and AKT1—proteins known to be central mediators of tumor progression and drug resistance.

Through this multi-dimensional computational strategy, we aimed to reveal the molecular basis underlying the therapeutic potential of Aloe vera in NSCLC, identify promising active compounds for further experimental validation, and provide theoretical support for the development of novel plant-derived anticancer agents with high efficacy and low toxicity.

## 2. Methods

### 2.1 Screening of Functional Components in Aloe vera

To systematically identify bioactive constituents of Aloe vera, we utilised the Traditional Chinese Medicine Systems Pharmacology Database and Analysis Platform (TCMSP; https://tcmsp-e.com), one of the most comprehensive repositories of phytochemicals and their predicted pharmacokinetic properties. “Aloe” was used as the search keyword, and candidate compounds were screened based on two widely adopted ADME parameters: (i) oral bioavailability (OB) ≥ 30%, which predicts the likelihood that a compound achieves effective systemic exposure following oral administration; and (ii) drug-likeness (DL) ≥ 0.18, an index that integrates structural and physicochemical similarity to known drugs. These criteria ensured the selection of compounds with sufficient drug-like behaviour for subsequent network and docking analysis.

### 2.2 Target Identification and Network Construction

For each shortlisted aloe compound, potential protein targets were extracted from TCMSP and further confirmed or supplemented by SwissTargetPrediction (STP; http://www.swisstargetprediction.ch) based on chemical structure similarity and machine-learning prediction models. The canonical SMILES strings of each compound were retrieved from PubChem to facilitate STP input. Only targets with non-zero prediction probability were retained. Protein names were standardised to gene symbols using UniProt (https://www.uniprot.org) with species restricted to Homo sapiens. Redundant entries from TCMSP and STP were merged to generate a unified compound–target dataset.

Disease-associated targets for NSCLC were collected from GeneCards (www.genecards.org) and OMIM (www.omim.org), using “non-small cell lung cancer” as the keyword. GeneCards relevance scores were used to prioritise high-confidence genes involved in NSCLC pathogenesis, signalling, or therapeutic response. The intersection between aloe-derived targets and NSCLC-related genes was obtained using the VENNY 2.1 (https://bioinfogp.cnb.csic.es/tools/venny/) tool to generate overlapping gene sets. These overlapping genes were imported into Cytoscape v3.10.3 to construct a compound–target–disease network, enabling visualisation of node connectivity and compound–gene associations.

### 2.3 Protein–Protein Interaction (PPI) Network and Topological Analysis

To evaluate functional associations among the overlapping targets, a high-confidence PPI network was generated using STRING (https://string-db.org/), with interaction confidence set to the highest threshold (>0.9). Unconnected nodes were removed to preserve biological relevance. The resulting network file was imported into Cytoscape for topological analysis.

Network centrality was assessed using the cytoNCA plugin, focusing on three key measures: Degree centrality (DC): number of direct connections, indicating regulatory prominence. Betweenness centrality (BC): frequency at which a node lies on shortest paths, reflecting information-flow control. Closeness centrality (CC): average distance to all other nodes, indicating signalling efficiency.

Targets with high centrality values typically play pivotal biological roles. In parallel, Hub genes were identified using the CytoHubba plugin based on Edge Percolated Component (EPC) scoring, which retains nodes resilient to edge removal. The Top 10 Hubba genes were considered core regulators in the aloe–NSCLC network.

### 2.4 GO and KEGG Enrichment Analysis

Functional enrichment analyses were performed using Metascape (https://metascape.org), which integrates multiple annotation databases including KEGG, GO, Reactome, BioGrid, and others. GO enrichment encompassed the domains of Biological Process (BP), Cellular Component (CC), and Molecular Function (MF). KEGG pathways were analysed to identify signalling cascades and disease pathways enriched among the overlapping targets. Statistical enrichment was based on a hypergeometric test, and significant terms were defined using a false discovery rate (FDR) < 0.05. The top 20 enriched GO terms and KEGG pathways were visualised using the WeBio platform (www.bioinformatics.com.cn). These analyses facilitated the identification of NSCLC-related biological functions, including proliferation, apoptosis, stress response, inflammation, kinase activity, and drug-resistance pathways.

### 2.5 Molecular Docking

Molecular docking was performed to predict the binding affinities and interaction modes between Aloe compounds and key NSCLC targets. Three-dimensional (3D) structures of the ligands were obtained from PubChem, whereas protein structures for p53, EGFR, EGFR T790M/C797S, and AKT1 were downloaded from the Protein Data Bank (PDB; www.rcsb.org). Redundant chains, water molecules, and crystallographic artefacts were removed, retaining biologically relevant monomers.

Docking was conducted using CB-Dock2 (http://clab.labshare.cn:10380/cb-dock2/php/blinddock.php) — a blind docking platform that automatically identifies optimal binding cavities and performs docking using AutoDock Vina scoring. Binding affinity was expressed as docking energy (kcal/mol), with values < −5.0 kcal/mol indicating strong spontaneous binding. The top-ranked poses were selected for visualisation, and 2D interaction diagrams, including hydrogen bonds, hydrophobic contacts, π-stacking, and salt bridges, were generated using Discovery Studio Visualizer.

### 2.6 Molecular Dynamics Simulation (MDS) and Binding Free Energy Analysis

To assess the dynamic stability of ligand–protein complexes, 100-ns all-atom molecular dynamics simulations were performed using Gromacs 2022. Protein topologies were constructed using the AMBER14SB force field, whereas ligand parameters were generated with the GAFF force field. Complexes were solvated in a TIP3P water box with appropriate counterions to achieve charge neutrality.

Energy minimisation was performed using the steepest descent method, followed by equilibration under NVT (100 ps) and NPT (100 ps) conditions at 298 K and 1 bar, respectively. Production runs lasted 100 ns with a 2-fs timestep. Electrostatic interactions were treated with the Particle-Mesh Ewald (PME) method, and non-bonded interactions were truncated at 1.2 nm. Hydrogen bonds were constrained with the LINCS algorithm. Trajectory analyses included: Root-mean-square deviation (RMSD) — evaluates structural stability. Root-mean-square fluctuation (RMSF) — assesses residue flexibility. Solvent-accessible surface area (SASA) — indicates ligand burial. Hydrogen bond count — reflects interaction strength. Radius of gyration (Rg) — measures compactness of the complex. Binding free energies were calculated using MMPBSA (g_mmpbsa), decomposing contributions of van der Waals, electrostatic, polar and nonpolar solvation energies.

## 3. Results

### 3.1 Network pharmacological analysis of Aloe–NSCLC targets

After retrieving and consolidating candidate compounds from TCMSP and corresponding predicted targets from TCMSP and SwissTargetPrediction, a total of 369 unique aloe-derived targets were identified. In parallel, interrogation of GeneCards and OMIM yielded 5519 NSCLC-associated targets, reflecting the extensive molecular heterogeneity and pathway complexity of NSCLC pathogenesis.

A Venn analysis revealed 268 overlapping targets between aloe constituents and NSCLC, representing a substantial intersection indicative of broad functional relevance (Fig. 1A). These overlapping genes were integrated into a compound–target–disease network using Cytoscape. As illustrated in (Fig. 1B), aloe bioactive molecules displayed extensive connectivity, each interacting with multiple targets. This network topology is characteristic of natural products, which often exert therapeutic effects through multi-target synergy rather than single-node inhibition.

**Figure 1.**
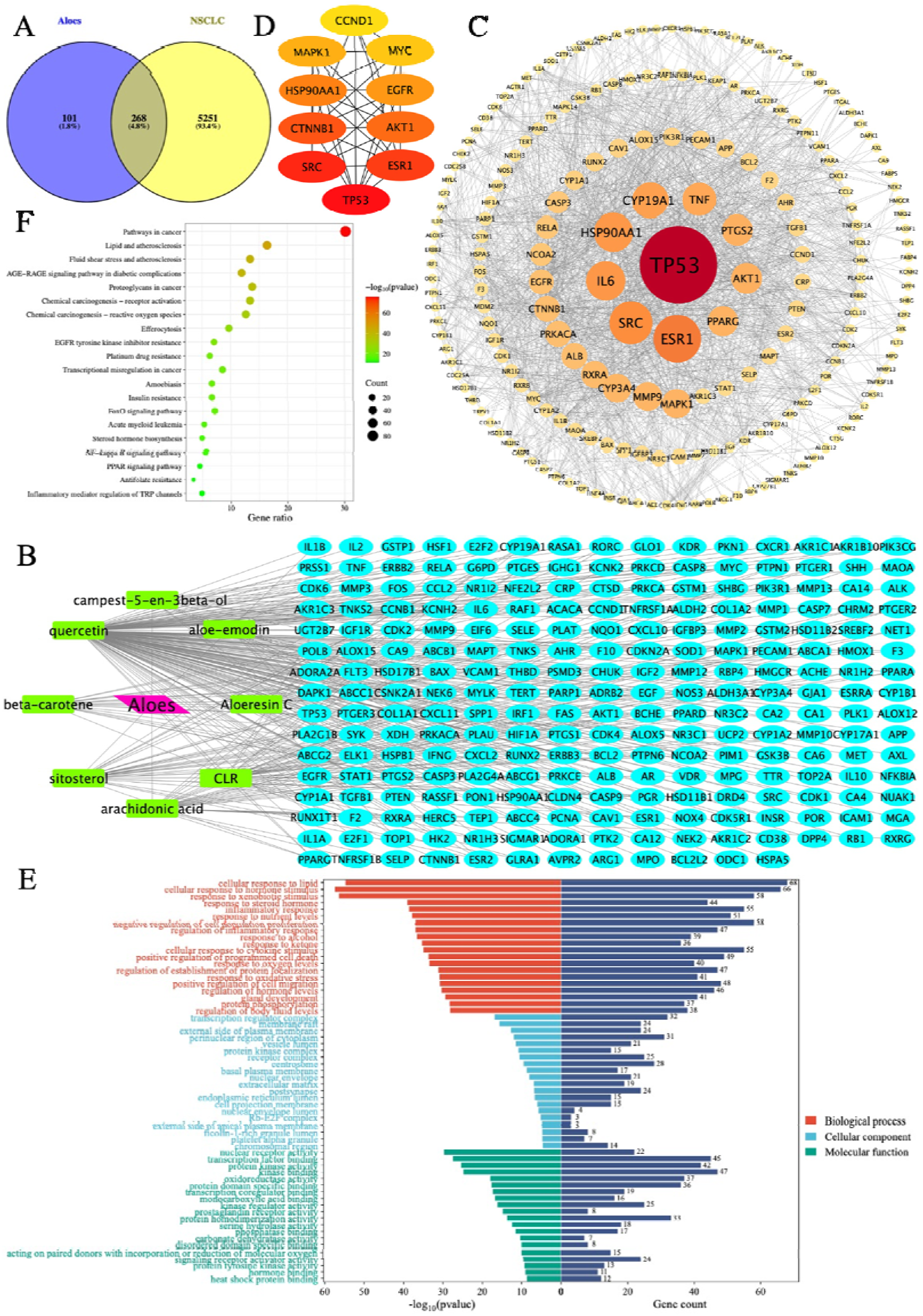
A: Venn diagram of aloe and NSCLC targets. Blue: targets of aloe yellow: targets of NSCLC. B: Aloes -NSCLC-Gene Network. Green: active ingredients of aloe blue: targets Gene Name. C: PPI analysis of targets. D: Top 10 hubba genes. E: GO enrichment analysis. F: KEGG Pathway Enrichment.

Inspection of the network revealed that compounds such as aloe-emodin, aloeresin C, quercetin, β-sitosterol, and campesterol derivatives exhibited particularly high degrees of connectivity. These polypharmacological features suggest that diverse aloe components may converge on common pathogenic pathways in NSCLC or complement one another through distributed signaling modulation.

To further elucidate the relational architecture among the overlapping targets, we constructed a protein–protein interaction (PPI) network using STRING (confidence > 0.9). After removal of isolated nodes, the resulting network demonstrated a tightly interconnected topology reflecting biological coordination among NSCLC-relevant pathways (Fig. 1C). Topological analysis using cytoNCA identified several central nodes based on BC, CC, and degree scores. TP53, EGFR, AKT1, STAT3, JUN, VEGFA, MAPK1, and TNF ranked among the highest, highlighting their essential roles in cellular proliferation, apoptosis, angiogenesis, inflammation, and drug resistance.

CytoHubba EPC analysis further confirmed TP53, AKT1, STAT3, EGFR, and VEGFA as top hub genes (Fig. 1D). TP53, which encodes the tumor suppressor p53, was the most prominent hub, consistent with its critical involvement in DNA damage response, apoptosis, genomic stability, and therapeutic resistance. Notably, EGFR and AKT1—a core receptor tyrosine kinase and a central effector of the PI3K/AKT pathway—also emerged among the strongest hubs, reflecting their pivotal role in NSCLC development and EGFR-TKI resistance.

Collectively, the network pharmacology results suggest that Aloe vera exerts anti-NSCLC effects through a dispersed but coordinated network of interactions targeting essential oncogenic pathways, potentially enabling multi-layered pathway inhibition and modulation of drug resistance mechanisms.

### 3.2 GO and KEGG enrichment analyses reveal Aloe–NSCLC pathway associations

To investigate the biological functions associated with the overlapping targets, GO and KEGG enrichment analyses were performed using Metascape. The top GO biological process (BP) categories enriched included response to oxidative stress, protein kinase activity regulation, cellular response to heterotrophic stimulus, apoptotic signaling modulation, DNA damage response, and Rb–E2F complex involvement (Fig. 1E). These categories directly align with hallmark hallmarks of NSCLC progression, including excessive proliferation, chronic inflammation, oxidative imbalance, and impaired apoptotic machinery.

In the cellular component (CC) category, enriched terms included nuclear chromatin, membrane raft, and cytosolic protein complex, consistent with the compartmentalised activation of signaling pathways such as EGFR, MAPK, and PI3K/AKT. Molecular function (MF) enrichment highlighted kinase binding, transcription factor binding, and enzyme regulatory activity, indicating that many aloe–NSCLC intersection genes encode nodes responsible for coordinating key intracellular signals.

KEGG pathway enrichment further underscored the involvement of critical NSCLC-associated pathways (Fig. 1F). The top enriched pathways included: EGFR tyrosine kinase inhibitor resistance; PI3K-AKT signaling pathway; p53 signaling pathway; MAPK signaling; TNF and IL-17 inflammatory pathways; Apoptosis; HIF-1 signaling; Pathways in cancer.

The inclusion of EGFR-TKI resistance among the top pathways provides direct computational evidence supporting the hypothesis that aloe-derived molecules may modulate or attenuate drug resistance mechanisms. Many of these pathways—particularly PI3K/AKT/mTOR and MAPK cascades—are known to be activated upon osimertinib or gefitinib failure, either through secondary EGFR mutations (T790M, C797S) or bypass pathway activation. Thus, the enrichment profile indicates that aloe components may exert therapeutic benefits not only by directly modulating tumor growth but also by suppressing resistance-driving pathways.

### 3.3 Molecular docking demonstrates strong binding of Aloe compounds with p53, EGFR, T790M/C797S, and AKT1

To identify key interaction pairs between aloe compounds and major NSCLC targets, molecular docking was performed on wild-type EGFR, EGFR T790M/C797S, AKT1, and p53. These proteins were selected due to their central involvement in tumor suppression (p53), oncogenic signaling (EGFR and AKT1), and resistance development (T790M and C797S mutations).

Docking results demonstrated that aloe-emodin, aloeresin C, quercetin, β-carotene, sitosterols, and campesterol derivatives all exhibited strong spontaneous binding to p53, with docking energies typically < −6.0 kcal/mol (Fig. 2). Many interactions involved hydrogen bonding to DNA-binding domain residues, hydrophobic stabilization within p53’s core scaffold, and π–π interactions with aromatic residues. These binding features suggest that aloe compounds could potentially influence p53 activation or stabilisation.

**Figure 2.**
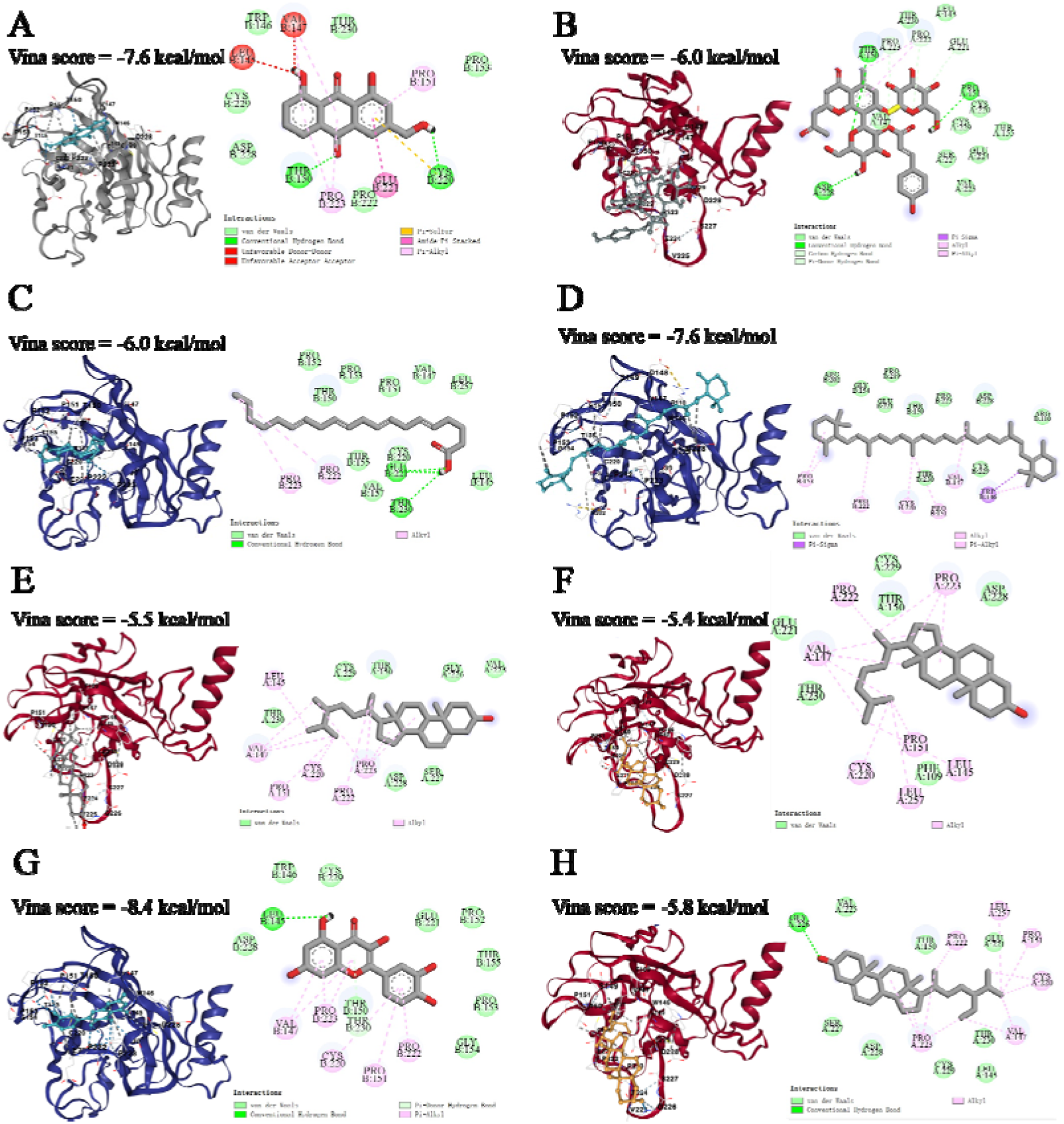
Molecular docking of p53 and aloe-emodin (A), aloeresin C (B), arachidonic-acid (C), beta-carotene (D), campest-5-en-3beta-ol (E), CLR (F), quercetin (G), sitosterol (H).

For wild-type EGFR, all eight compounds displayed favourable binding conformations (Fig. 3), often occupying the ATP-binding cleft or allosteric regulatory pockets. Hydrophobic scaffolds such as β-sitosterol and CLR provided substantial nonpolar burial, while polyphenolic compounds such as quercetin engaged in hydrogen bonding with hinge-region residues exemplified by Met793.

**Figure 3.**
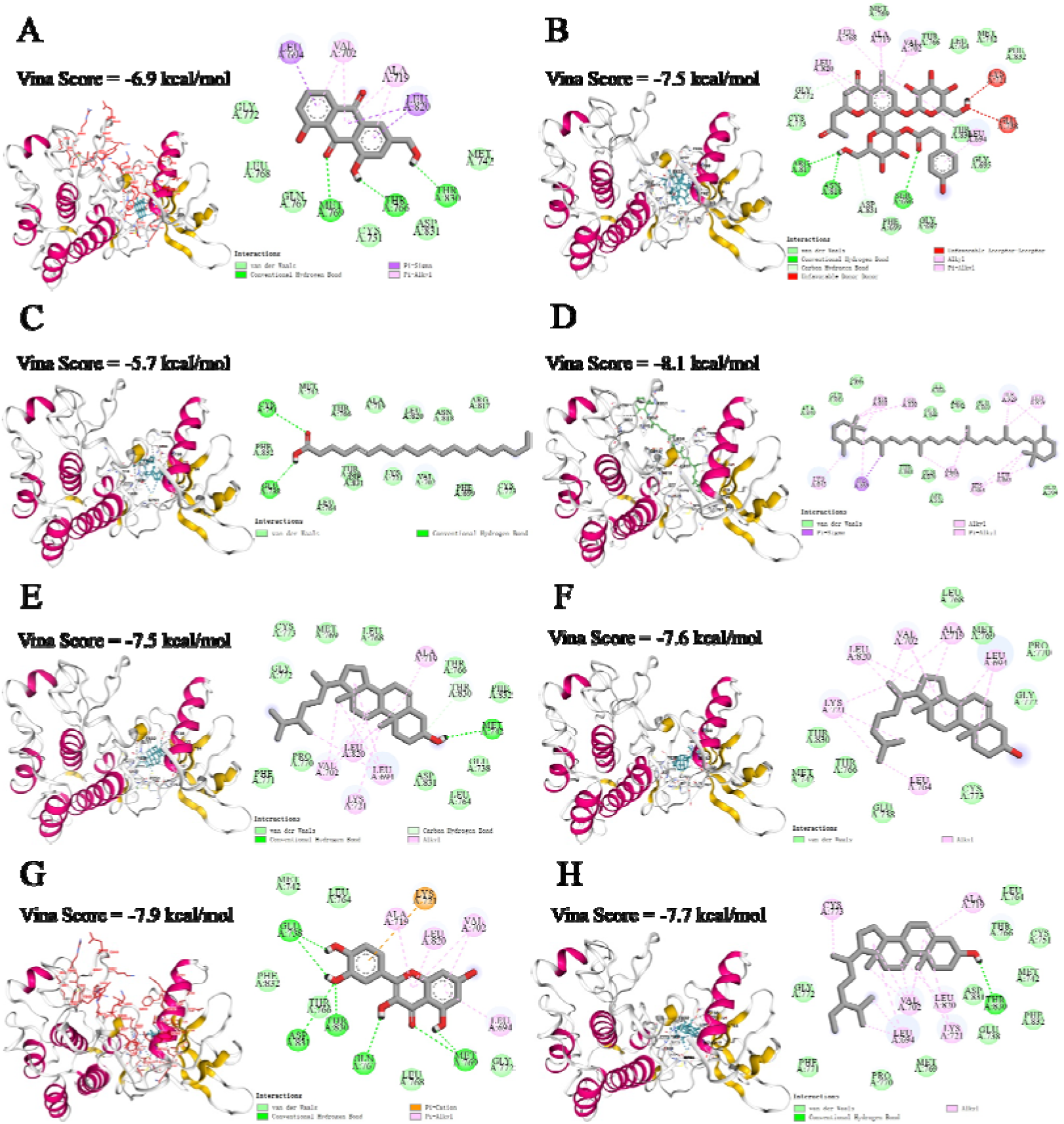
Molecular docking of EGFR and aloe-emodin (A), aloeresin C (B), arachidonic-acid (C), beta-carotene (D), campest-5-en-3beta-ol(E), CLR (F), quercetin (G), sitosterol (H).

Strikingly, docking against the drug-resistant EGFR T790M/C797S mutant revealed robust binding for all compounds except β-carotene (Fig. 4). Aloeresin C displayed the strongest binding affinity (−8.2 kcal/mol), forming multiple hydrogen bonds and hydrophobic contacts near the mutated residues. This suggests that certain aloe constituents may maintain affinity for EGFR even in its TKI-resistant conformations.

**Figure 4.**
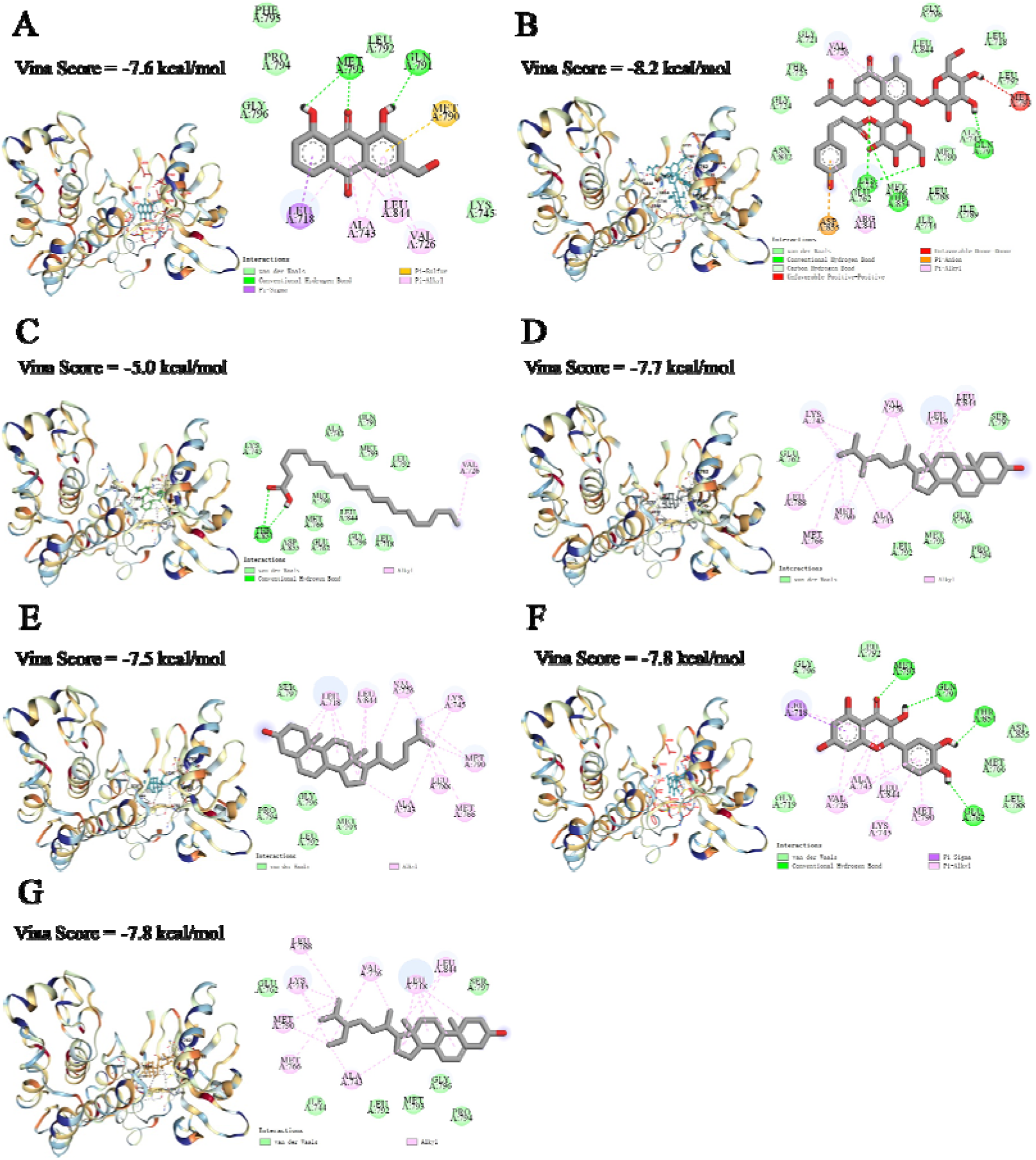
Molecular docking of T790M/C797S and aloe-emodin (A), aloeresin C (B), arachidonic-acid (C), campest-5-en-3beta-ol (D), CLR (E), quercetin (F), sitosterol (G).

Docking with AKT1 revealed consistent moderate-to-strong binding across seven compounds (Fig. 5). Ligands typically interacted with residues surrounding the pleckstrin homology (PH) domain or catalytic domain, indicating possible inhibition of AKT1 activation or substrate engagement. Given AKT1’s central role in the PI3K/AKT/mTOR axis, this provides further support for aloe-mediated pathway suppression. Overall, the molecular docking data highlight aloe compounds as promising ligands capable of targeting multiple nodes of NSCLC signaling networks, including those central to therapeutic resistance.

**Figure 5.**
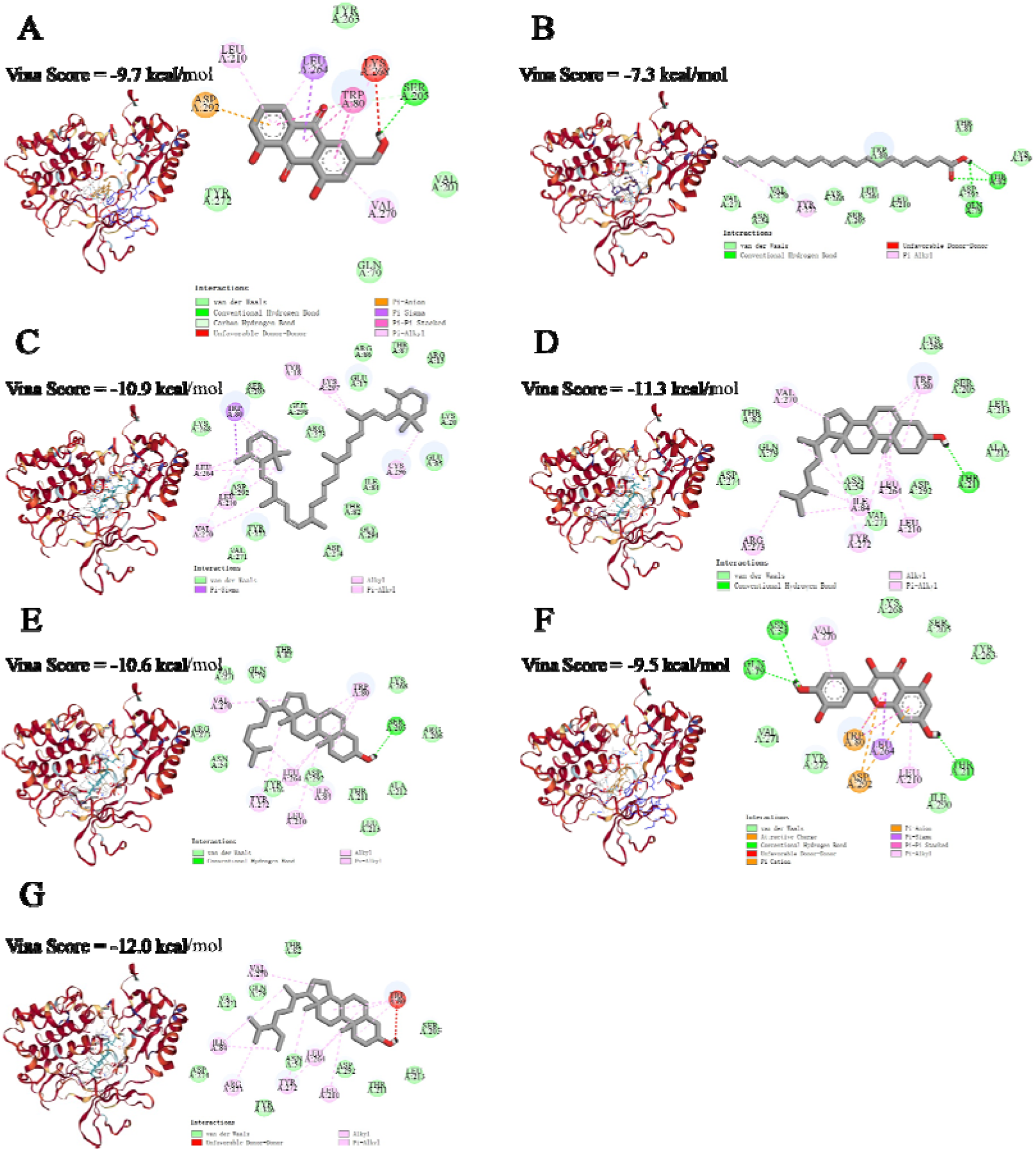
Molecular docking of AKT1 and aloe-emodin (A), arachidonic-acid (B), beta-carotene (C), campest-5-en-3beta-ol (D), CLR (E), quercetin (F), sitosterol (G).

### 3.4 Molecular dynamics simulation validates ligand–protein stability and favourable energy landscapes

To evaluate whether aloe compounds maintain stable interactions with target proteins under physiological conditions, 100-ns molecular dynamics simulations were performed. The stability of complexes was examined using RMSD, RMSF, SASA, hydrogen bond counts, radius of gyration, conformational overlays, electrostatic potential mapping, and free energy landscape (FEL) analysis. RMSD trajectories demonstrated that protein–ligand complexes gradually stabilised after initial adaptation periods (Fig. 6A). Complexes consistently reached equilibrium within 20–30 ns, with fluctuations typically < 0.25 nm, indicating robust structural stability. Ligands remained anchored within binding pockets throughout the simulation. RMSF analysis revealed decreased flexibility in residues surrounding ligand-binding cavities compared to apo proteins (Fig. 6B), suggesting ligand-induced local stabilisation. This effect was particularly pronounced for EGFR and AKT1 complexes. SASA profiles demonstrated progressive burial of ligands into hydrophobic pockets (Fig. 6C), supporting stable encapsulation. The solvent-exposed area consistently decreased over time. Hydrogen bond analysis indicated that complexes maintained 2–7 hydrogen bonds on average (Fig. 6D), reflecting moderate-to-strong polar interactions that contribute to binding stability. Radius of gyration (Rg) exhibited a decreasing trend for most complexes (Fig. 6E), suggesting enhanced structural compactness and favourable folding states upon ligand binding. Overlay of simulated conformations showed minimal deviation from initial docked structures (Fig. 6F), reinforcing the structural integrity of the complexes. Electrostatic potential mapping (Fig. 6G) indicated that ligand binding is facilitated by favourable electrostatic complementarities, including interactions between polar functional groups and charged residues within the binding cavity. Finally, free energy landscape (FEL) analysis revealed that complexes converged into a single low-energy basin (Fig. 6H), indicative of stable thermodynamic states and minimal conformational drift. Together, MDS results support the notion that aloe compounds form stable and energetically favourable interactions with key NSCLC targets, providing atomistic evidence of their inhibitory potential.

**Figure 6.**
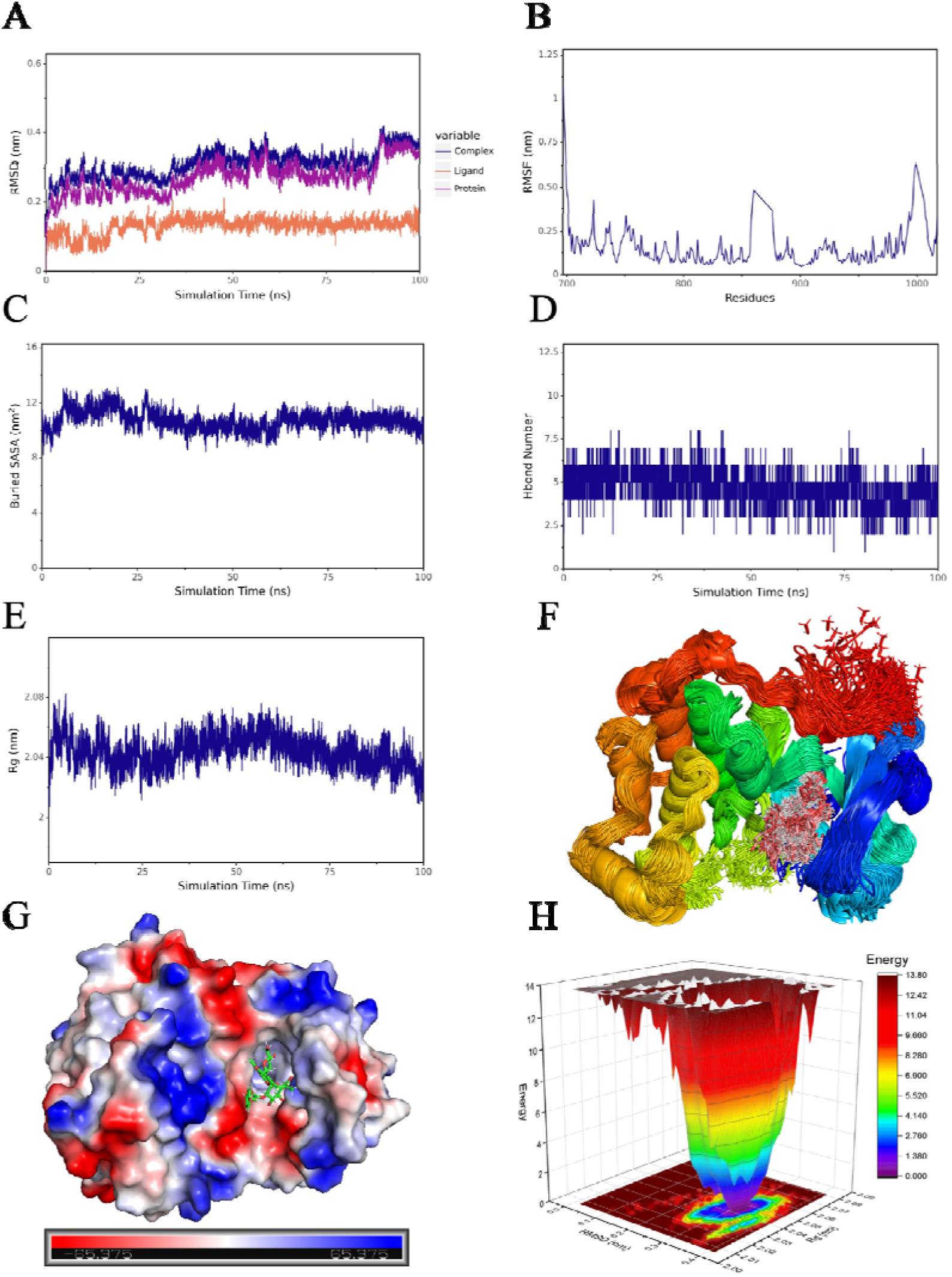
A: RMSD of complexes, proteins, and small molecule. B: RMSF of proteins in complexes. C: SASA between small molecules and proteins. D: Hbond number. E: Rg of the complex. F: Simulated conformation overlay. G: Surface electrostatic potential of small molecule binding proteins (unit: kcal/mol). H: Free energy landscape.

## 4. Discussion

Aloe is distributed extensively across Asia, Africa, and other regions, and has been utilised for the treatment of illnesses, beauty care, and health maintenance for millennia. Due to the rich compounds in its leaves, consuming the juice orally or applying it topically offers numerous health benefits [23]. As a perennial herbaceous plant belonging to the Asphodelaceae family within the Asparagales order, aloe exhibits a high level of safety for vital organs such as the liver, kidneys, and lungs, making it a suitable candidate for pharmaceutical research and development [24]. In previous studies, aloe vera has been applied in antiviral therapy [25], alleviating photoaging of the skin [26], and preventing radiation-induced dermatitis [27]. In recent years, aloe vera has also demonstrated significant potential value in cancer treatment, with colon cancer being the most extensively studied. The active components of aloe induce mitochondrial phagocytosis in colon cancer cells, leading to cancer cell death [19, 28], They also inhibit cell migration, survival, and stemness in colorectal cancer [20]. However, research on the use of aloe vera for combating lung cancer and drug resistance remains limited, and the underlying mechanisms are not yet fully understood [21]. Network pharmacology is an online, efficient, and low-cost approach to drug discovery that has gained widespread popularity among scholars in recent years. This study employs network pharmacology methods to comprehensively investigate the mechanism by which aloe vera combats non-small cell lung cancer. The human tumour suppressor gene TP53 is closely associated with the cell cycle and genomic stability, encoding the p53 protein. In the event of cells receiving stress signals, such as DNA damage, there is a dramatic surge in p53 levels. As a transcription factor, p53 has been shown to bind to specific DNA sequences in order to regulate gene expression, thus rendering it one of the key targets in NSCLC [29]. In both the PPI network (Fig 1C) and TOP10 Hubba gene (Fig 1D), TP53 occupies a central position. The molecular docking results indicate that eight active components in aloe vera can spontaneously form strong bonds with p53, consistent with previous research findings [30]. These molecules have been observed to bind directly to p53, thereby either inhibiting cancer cell proliferation or inducing cancer cell apoptosis.

Lung cancer is a prevalent form of cancer with a high mortality rate, with NSCLC accounting for over 80% of all lung cancers. EGFR is a transmembrane protein that transmits growth factor signals, possessing cytoplasmic kinase activity, with its extracellular receptor domain binding to EGF [10]. In NSCLC, EGFR is a key target and hallmark protein, exhibiting abnormally elevated expression levels. EGFR tyrosine kinase inhibitors (TKIs) are a frequently employed class of pharmaceutical agents in the management of NSCLC. First-generation EGFR-TKIs (gefitinib, erlotinib) reversibly bind to the epidermal growth factor receptor (EGFR), resulting in suboptimal therapeutic outcomes. It has been established that patients develop the T790M resistance mutation after approximately one year of treatment. Second-generation EGFR-TKIs such as afatinib and dacomitinib, irreversibly bind to the EGFR itself. However, their capacity to inhibit multiple targets results in a greater propensity for adverse effects and an inability to circumvent drug resistance. Third-generation EGFR-TKIs (osimertinib, amelanotib) were developed to address T790M resistance, but they can induce new C797S resistance mutations [31]. As demonstrated in preceding studies, approximately 5% of patients with activating EGFR mutations in NSCLC develop resistance to EGFR-TKIs through SCLC transformation during treatment [11, 12]. In the context of KEGG pathway enrichment analysis (Fig 1F), the EGFR-TKI resistance pathway was found to be enriched. The wild-type EGFR and T790M/C797S mutations were selected for molecular docking with the active components of aloe vera. The results indicate that all eight compounds exhibit strong binding to the EGFR (Fig 2). With the exception of β-carotene, all compounds exhibited strong binding to T790M/C797S (Fig 3). Aloeresin C exhibits the lowest binding energy with T790M/C797S (-8.2 kcal/mol) and forms hydrogen bonds and other high-energy bonds (Fig 3B). MDS results indicate that the two are stably combined.

Abnormal activation of the PI3K/AKT/mTOR pathway is a prerequisite for NSCLC progression and contributes to the development of resistance to EGFR-TKIs [32]. AKT1 is a core component of the PI3K/AKT/mTOR pathway and plays a crucial role in cancer cell proliferation and migration [33]. The AKT1 protein has been shown to act as a "cell survival switch". Activation of this pathway is initiated by upstream signals, such as those derived from the EGFR, which subsequently trigger the transmission of signals to the cell, thereby inducing uninterrupted proliferation through the process of phosphorylation at the molecular level [34]. AKT1 demonstrates increased levels of activity in EGFR-TKI-resistant cells in comparison to NSCLC parent cells. It has been demonstrated in previous studies that the elimination of MEK1 and AKT1/2 results in the inhibition of the growth of osimertinib-resistant cells and the partial restoration of osimertinib sensitivity [35].

Molecular docking results indicate that, with the exception of aloeresin C, all seven remaining aloe components exhibit strong binding affinity with AKT1 (Fig 4). This finding indicates that the active components present in aloe vera may directly bind to AKT1, thereby inhibiting its reception of upstream signals and its transmission of downstream signals that drive the cancer cell cycle.

In summary, a network pharmacology research strategy integrating molecular docking and MDS was employed to systematically reveal the potential multi-target, multi-pathway mechanism of aloe vera’s primary bioactive components against NSCLC. The potential applications of aloe vera in the field of cancer treatment warrant further exploration. Subsequent research should prioritise validating core predicted targets and pathways, and translating computational data into conclusive biological evidence. Multi-omics technologies, including metabolomics and proteomics, facilitate a more comprehensive mapping of the dynamic interactions of aloe vera components within NSCLC cells. The integration of multi-omics technologies constitutes a fundamental building block, providing a robust framework for subsequent exploration and advancement in this field. Concurrently, the exploration of combination therapy regimens that integrate aloe vera’s active components with existing chemotherapy drugs or targeted therapies to assess whether they can enhance efficacy or mitigate side effects holds significant clinical and translational implications. This progressive research strategy, which involves transitioning from prediction to validation and from the big picture to the details, will unlock new medical value for the traditional Chinese medicinal plant aloe vera and propel breakthroughs in this field.

### 4.8 Conclusions

This study provides the first systems-level mechanistic exploration of Aloe vera’s anti-NSCLC potential, identifying a multi-target, multi-pathway mode of action that includes p53 activation, EGFR inhibition (including resistant mutations), AKT1 suppression, and modulation of inflammatory and apoptotic pathways. Aloe’s bioactive molecules demonstrate strong binding affinities and dynamic stability with key NSCLC targets, highlighting their potential as adjunctive or innovative therapeutic agents. These findings lay a robust theoretical and computational foundation for future experimental validation and the development of natural product–derived anticancer drugs with low toxicity and broad molecular coverage.

## Acknowledgements

This research was supported by the National Key R&D Program of China [Project No.: 2022YFD22006].

